# Optogenetic fMRI reveals therapeutic circuits of subthalamic nucleus deep brain stimulation

**DOI:** 10.1101/2024.02.22.581627

**Authors:** Yuhui Li, Sung-Ho Lee, Chunxiu Yu, Li-Ming Hsu, Tzu-Wen W. Wang, Khoa Do, Hyeon-Joong Kim, Yen-Yu Ian Shih, Warren M. Grill

**Author notes:** These authors contributed equally (Yuhui Li, Sung-Ho Lee). Correspondence for Warren M. Grill, for Yen-Yu Ian Shih.

## Abstract

While deep brain stimulation (DBS) is widely employed for managing motor symptoms in Parkinson’s disease (PD), its exact circuit mechanisms remain controversial. To identify the neural targets affected by therapeutic DBS in PD, we analyzed DBS-evoked whole brain activity in female hemi-parkinsonian rats using function magnetic resonance imaging (fMRI). We delivered subthalamic nucleus (STN) DBS at various stimulation pulse repetition rates using optogenetics, allowing unbiased examinations of cell-type specific STN feed-forward neural activity. Unilateral STN optogenetic stimulation elicited pulse repetition rate-dependent alterations of blood-oxygenation-level-dependent (BOLD) signals in SNr (substantia nigra pars reticulata), GP (globus pallidus), and CPu (caudate putamen). Notably, these manipulations effectively ameliorated pathological circling behavior in animals expressing the kinetically faster Chronos opsin, but not in animals expressing ChR2. Furthermore, mediation analysis revealed that the pulse repetition rate-dependent behavioral rescue was significantly mediated by optogenetically induced activity changes in GP and CPu, but not in SNr. This suggests that the activation of GP and CPu are critically involved in the therapeutic mechanisms of STN DBS.

## INTRODUCTION

Deep brain stimulation (DBS) is a well-established neurosurgical therapy for controlling motor symptoms in humans with Parkinson’s disease (PD) who are unresponsive to dopaminergic medications or have developed levodopa-induced motor complications. DBS for PD involves implanting stimulating electrodes in subcortical brain areas, including subthalamic nucleus (STN). STN DBS decreases motor disability and enhances quality of life ^1^, and may delay symptom progression in PD patients ^2,3^.

Despite the clear clinical benefits of STN DBS in PD, its underlying neural mechanisms remain elusive. One obstacle is that electrical stimulation non-specifically activates both local neurons and axons around the stimulation site, which in turn alters neural activity across widespread areas of the brain ^4–8^. Accordingly, several hypotheses have been proposed for the neural pathways involved for therapeutic effects of STN DBS. The utilization of optogenetics offers a promising approach to activate specific cell types ^9^, which facilitates circuit dissection of the therapeutic STN DBS mechanisms in PD. Notably, several pioneering studies using Channel Rhodopsin-2 (ChR2) have reported that STN DBS can achieve its therapeutic effects by antidromic activation of motor cortical neurons via the hyper-direct pathway ^10,11^, shifting the focus of DBS research toward this pathway, rather than other STN feed-forward circuits. However, it is worth noting that the widely used ChR2 suffers from slow kinetics and cannot to follow photostimulation above 100 Hz ^12,13^. Considering that high pulse repetition rate stimulation (>100 Hz) is a hallmark of therapeutic STN DBS in PD, it is likely that the contributions of circuits downstream of STN have been overlooked. Indeed, a recent study reported robust behavioral therapeutic effects by stimulating STN cell bodies without direct engagement of its motor afferents ^14^. Despite this critical finding, it remains unclear how activity changes in different brain regions downstream of STN DBS are causally related to the behavioral rescue in PD. The potential for STN DBS to produce activation or inhibition in various related, yet anatomically distinct brain regions makes functional magnetic resonance imaging (fMRI) an ideal tool to decipher the circuit mechanisms governing therapeutic STN DBS. fMRI enables mapping whole-brain evoked responses in an unbiased manner, and such brain-wide patterns of activity changes during different stimulation parameters represent critical information to reveal the precise neural circuit changes that are necessary for therapeutic STN DBS ^15–21^.

Here, we combined optogenetic activation of STN neurons and fMRI to map the brain areas that are modulated by stimulating STN cell bodies in hemi-parkinsonian rats. We varied the pulse repetition rate, which is known to modulate the therapeutic effects of DBS ^22,23^, and we hypothesized that brain areas mediating the effects of STN DBS would exhibit responses differentially tuned to the pulse repetition rate in a manner correlated to the behavioral effects. We used an ultrafast opsin (Chronos)^24^ as required to follow faithfully the high pulse repetition rates necessary for effective STN DBS ^14^. We found that a range of subcortical brain regions, including substantia nigra pars reticulata (SNr), globus pallidus (GP), and caudate putamen (CPu) were activated by STN optogenetic DBS, and that the activation of GP and CPu causally mediate the pulse repetition rate-dependent therapeutic effect of STN DBS in PD.

## RESULTS

### Histological validation of the dopaminergic degeneration, virus expression and optical fiber implants

To evaluate DBS pulse repetition rate-dependent effects on the complex STN-related circuitry, we conducted a series of optogenetic-fMRI experiments in hemi-parkinsonian rats. For each optogenetic-fMRI experiment, blue light (473 nm) pulse-trains were delivered by fiber optic to the STN to activate either Chronos or ChR2 fused with green fluorescent protein (GFP) expressed by an adeno-associated-virus (AAV) vector. Expression was directed to excitatory neurons in STN under control of the calmodulin kinase IIα (CaMKIIα) promoter. Animals used in behavioral and imaging data collection (Chronos group, n = 5; ChR2 group, n = 4, **Figure 1A**) exhibited degeneration of dopaminergic cells in substantia nigra pars compacta (SNc) in the hemisphere receiving 6-hydroxydopamine (6-OHDA) injection (**Figure 1B**), as indicated by the clear reductions in tyrosine hydroxylase (TH) immunoreactivity (**Figure 1C**). Expression of Chronos or ChR2 in STN was confirmed by GFP (**Figure 1D**), and the locations of implanted optical fibers in STN were confirmed by structural MRI (**Figure 1E-F**).

**Figure 1.**
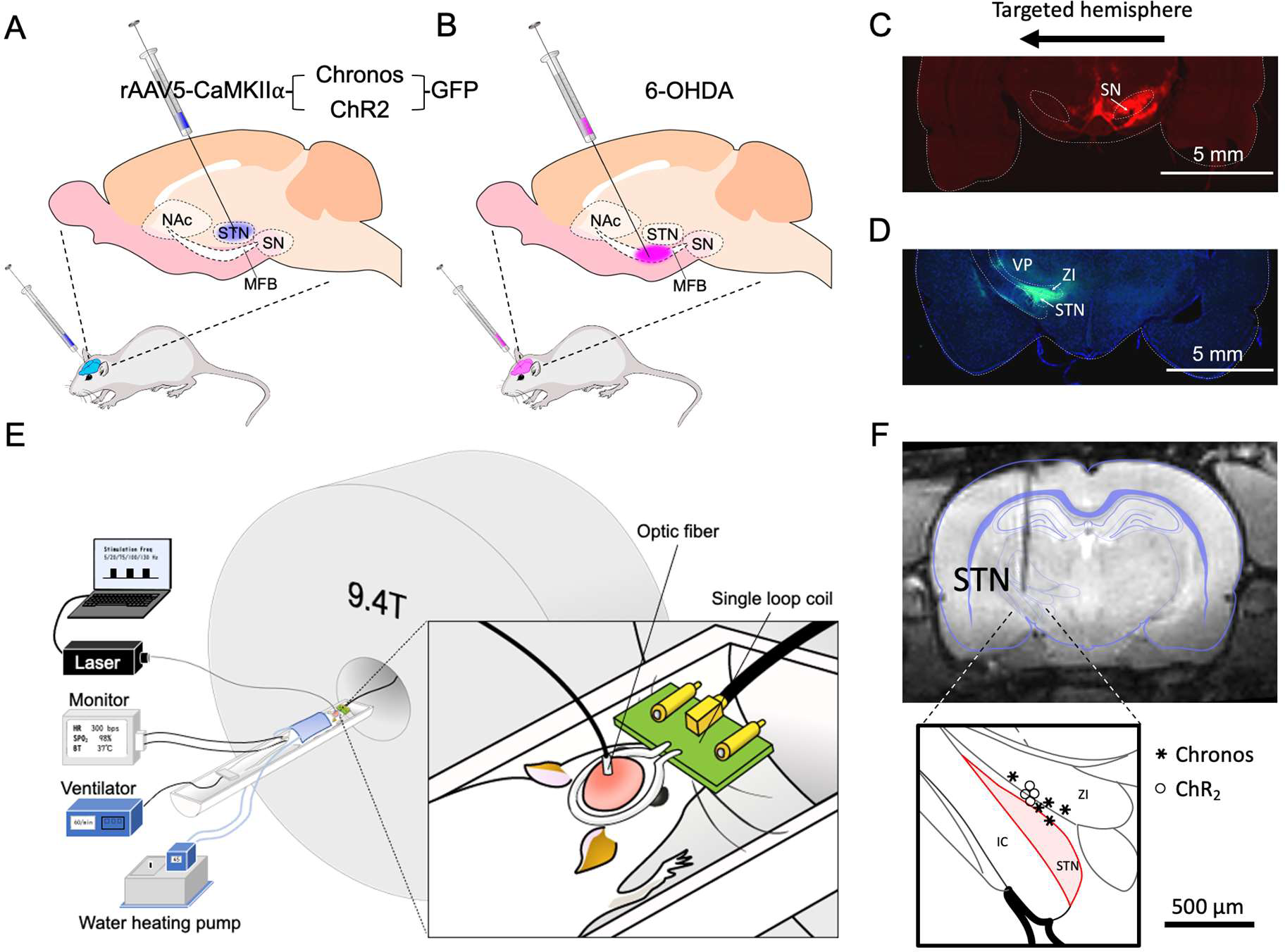
Chronos or ChR2 expression, TH immunofluorescence, and optical fiber implant location in hemi-parkinsonian rats. (A-B) schematic illustration of virus (A), and 6-OHDA (B) injections. (C) TH immunofluorescence showing 6-OHDA lesion of dopaminergic neurons in the left substantia nigra (SN). (D) Fluorescent micrograph showing expression of Chronos-GFP in the left STN. (E) Schematic illustration of MRI experimental setup. (F) Anatomical MR image showing the optical fiber track in the brain and location of the tip of the optical fibers in STN across all animals used in this study. The optical fiber implantation was targeted 300 µm above the STN viral expression site, allowing illumination of the STN.

### STN optogenetic stimulation reduced pathological motor behavior

The motor dysfunction induced by 6-OHDA lesion was characterized behaviorally by ipsilateral circling (**Figure 2A**) following administration of methamphetamine (mean ± SD = 8.85 ± 5.87 turns/min, n = 9; p = 0.0039, Mann-Whitney test compared to zero). Pathological circling behavior was ameliorated by 130 Hz optogenetic STN DBS in the Chronos group (F_(2, 75)_ = 73.4, p < 0.001, one-way repeated measure ANOVA), but not in the ChR2 group (F_(2, 60)_ = 1.52, p = 0.23). The effects of optogenetic DBS on angular velocity was dependent on pulse repetition rate (**Figure 2B-D**; two-way ANOVA with significant main effect of the pulse repetition rate (F_(4, 260)_ = 6.77, p < 0.001) and opsin type (F_(1, 260)_ = 78.6, p < 0.001), and a significant interaction between factors (F_(4, 260)_ = 7.96, p < 0.001). *Post hoc* paired comparisons showed that high pulse repetition rate optogenetic DBS (75, 100 and 130 Hz) applied to animals in the Chronos group reduced ipsilateral circling (all p < 0.01), whereas no changes in circling rate were detected during low rate optogenetic DBS (5 and 20 Hz) in the Chronos group or during optogenetic DBS at any pulse repetition rate in the ChR2 group (all p > 0.1). Importantly, neither optogenetic pulse repetition rate nor opsin type affected the linear speed of animals’ motion (**Figure 2E**; two-way ANOVA with non-significant main effects of pulse repetition rate (F_(4, 260)_ = 1.03, p = 0.39) or opsin type (F_(1, 260)_ = 2.36, p = 0.13), and a non-significant interaction between factors (F_(4, 260)_ = 1.42, p = 0.23). These results are consistent with previous studies of the effects of electrical ^25^ and optogenetic DBS ^14^, and established the comparative effectiveness of different rates and different optogenetic constructs prior to fMRI studies of the effects of optogenetic STN stimulation.

**Figure 2.**
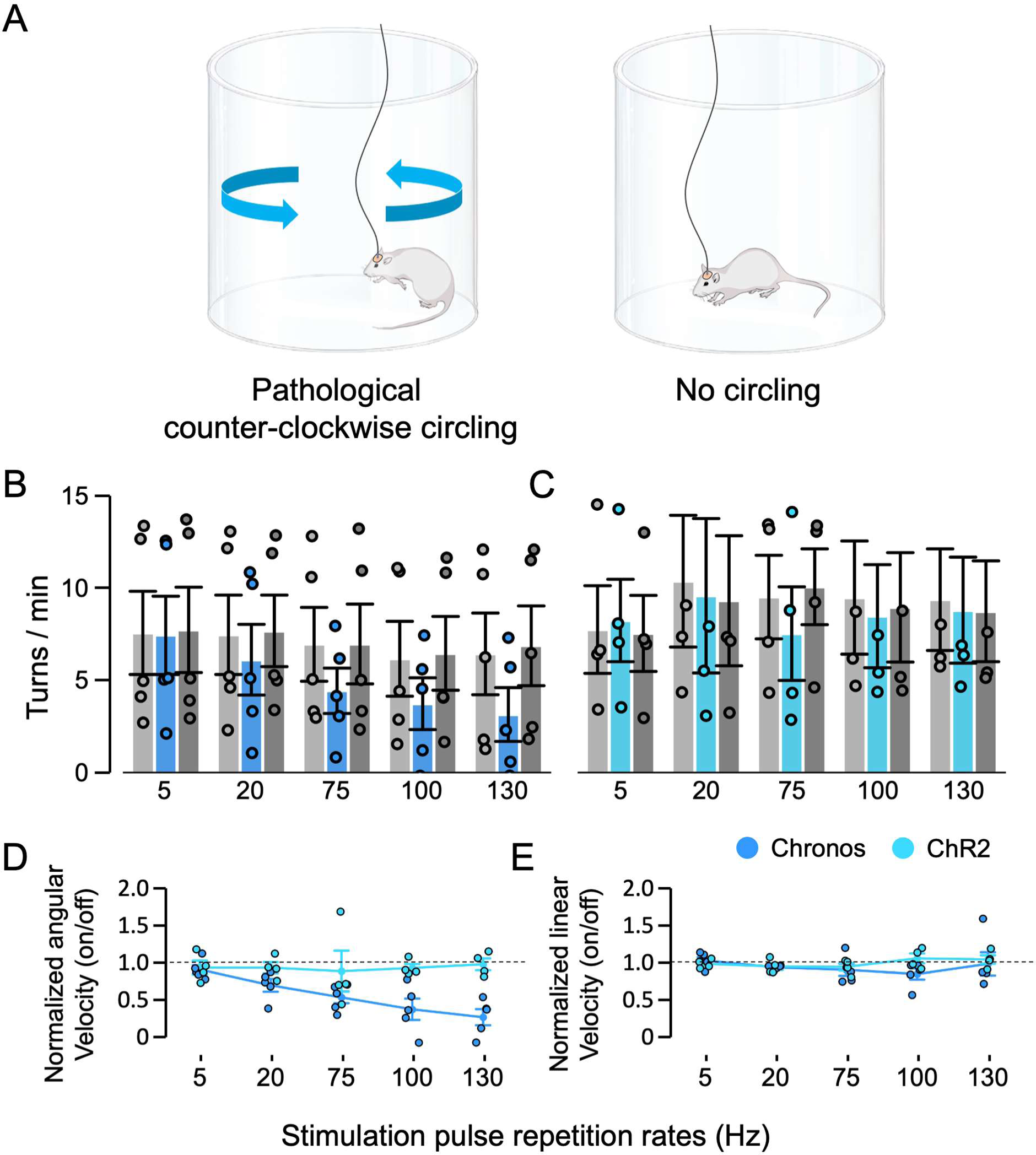
Effects of STN optogenetic DBS at different pulse repetition rates on the angular and linear velocities in methamphetamine-induced circling test. (A) Schematic illustrations of the pathological (counter-clockwise) circling behavior and non-pathological behaviors (no circling). (B-C) Measured number of turns per minute in Chronos (n=5) and ChR2 (n=4) rats. (D) Normalized angular velocity and (E) normalized linear velocities during stimulation in two groups of rats. STN optogenetic DBS pulse repetition rate has a significant effect on angular velocity in Chronos rats but not in ChR2 rats. DBS had no effect on linear velocity in either group of rats. Data are expressed as mean ± sem.

### *STN optogenetics DBS* activated three nuclei: *SNr, GP and CPu* within the basal ganglia

Next, the same cohorts of Chronos and ChR2 rats underwent fMRI. Six randomly ordered paradigms were used (5 different pulse repetition rate conditions plus no stimulation condition, **Figure 3A**), resulting in a group of n = 90 epochs for Chronos rats and n = 72 epochs for ChR2 rats per paradigm. Together, these experiments provided a grand total of n = 540 and 432 fMRI epochs for Chronos and ChR2, respectively, for the entire study. A general linear model (GLM) analysis was conducted to evaluate voxel-wise response patterns of unilateral STN optogenetic DBS evoked blood-oxygenation-level-dependent (BOLD) fMRI signal changes in hemi-parkinsonian rats. The group-averaged response maps showing each stimulation pulse repetition rate are presented by the type of opsin in **Supplementary Figure S1**, and reveal that certain brain regions responded differently during stimulation with low pulse repetition rates (5 and 20 Hz), which correspond to behaviorally ineffective STN optogenetic DBS, versus during stimulation with high pulse repetition rates (above 75 Hz), which are associated with behaviorally effective optogenetic DBS. In the Chronos group, within-group comparisons of high (75, 100, and 130Hz) and low (5 and 20Hz) pulse repetition rate ranges revealed significantly enhanced positive BOLD responses in ipsilateral SNr, SC (superior colliculus), GP, LHb (lateral habenula nucleus) and PTg (pedunculotegmental nucleus) during high-pulse repetition rate DBS (**Supplementary Figure S2**). In the ChR2 group, same analyses revealed significantly enhanced positive BOLD responses in ipsilateral medial forebrain bundle (MFB) and enhanced negative BOLD responses in ipsilateral CPu and contralateral SC (**Supplementary Figure S2**). Collectively, there were clear differences in the modulation of STN feedforward activity mediated by the behaviorally-effective Chronos construct and the behaviorally ineffective and kinetically limited ChR2 opsin.

**Figure 3.**
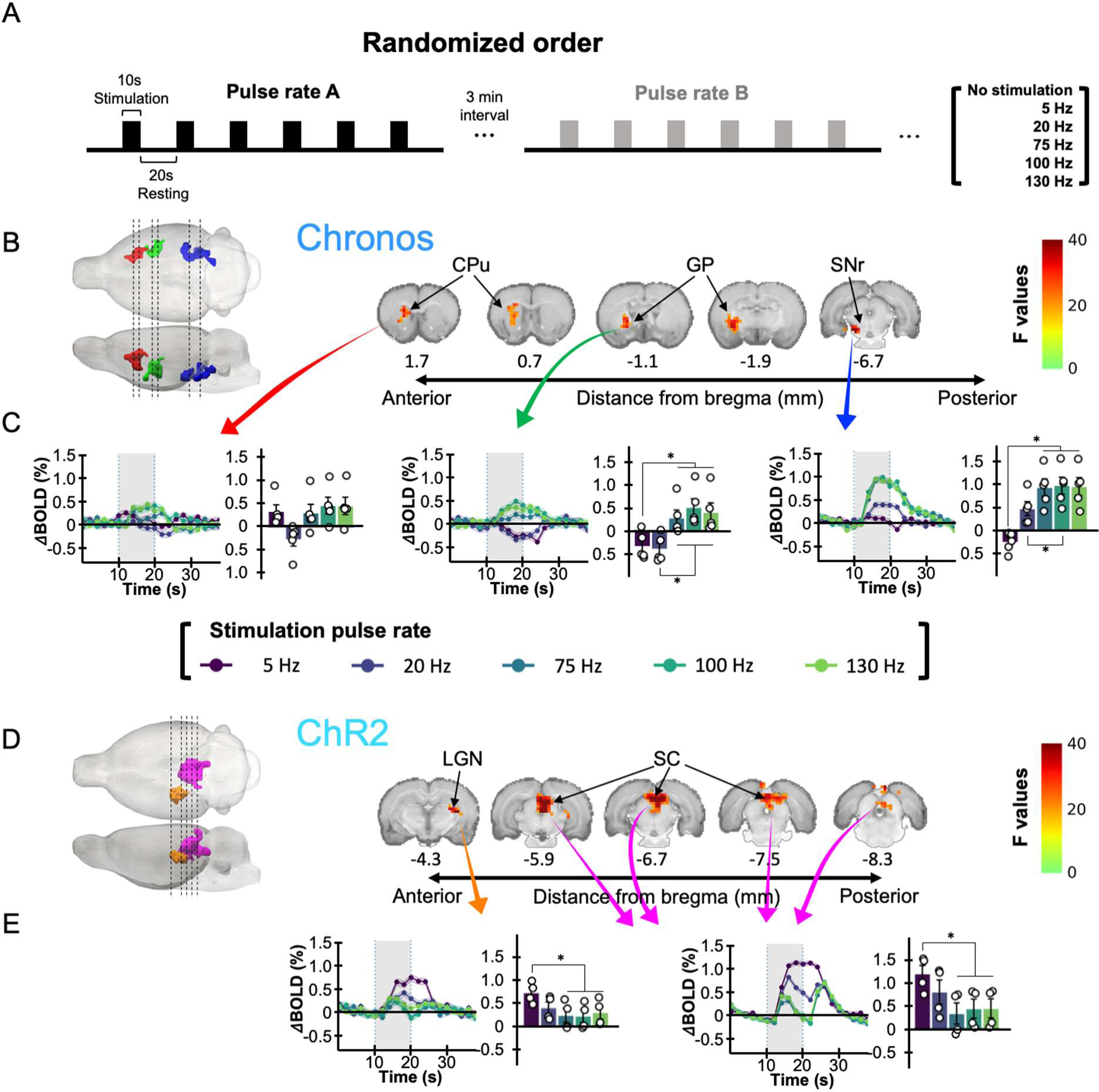
Brain-wide fMRI responses to STN optogenetic DBS. (A) Schematic illustration of the study design, including stimulation paradigms. (B-E) Voxel-wise analysis followed by *post ho*c tests depict significant pulse repetition rate-dependent BOLD responses during optogenetic STN DBS. The 3D representations on the left display the significant clusters for Chronos (B) and ChR2 (D), with statistical parametric maps shown on the right. The BOLD response patterns from respective cluster ROIs for the Chronos and ChR2 groups are shown in (C) and (E), respectively. The stimulation epoch is indicated by the gray-shaded band. Solid lines and color shadings of the time-courses represent mean ± sem. All bar graphs extract the peak of BOLD percentage changes during the stimulation period. * denotes p < 0.05 by T-test.

We then utilized a linear mixed effect regression model coupled with ANOVA to assess the voxel-wise effects of stimulation pulse repetition rate, opsin type, and their interaction on the BOLD signal using the data including both Chronos and ChR2 groups. This analysis revealed the main effect of pulse repetition rate across five spatially distinct clusters, depicting the common effects seen in both subject groups (**Supplementary Figure S3**). We also observed a significant interaction between the pulse repetition rate and opsin type in two additional clusters (**Supplementary Figure S4**), suggesting the choice of opsin is critical for fMRI outcomes, as it was for behavioral effects. It is important to highlight that the main effect of pulse repetition rate refer to common effect shared by both groups. Subsequent analyses focused on examining the effect of stimulation pulse repetition rate within each group to elucidate their distinct patterns. Considering the predominant opsin effects, we further conducted voxel-wise analyses to explore the pulse repetition rate effects within each individual subject group. In Chronos group, we revealed three brain regions, namely, ipsilateral CPu, GP and SNr (**Figure 3B**), that were significantly modulated by the stimulation pulse repetition rate. Analysis of time-courses in these regions in the Chronos group showed positive BOLD responses during 75-130Hz stimulation and negative or weak-positive BOLD responses during 5-20 Hz stimulation (**Figure 3C**). Notably, BOLD responses in these regions in the ChR2 group did not exhibit a significant increase at 75Hz and above, in contrast to the Chronos group (**Supplementary Figure S5**). Interestingly, the BOLD responses in GP and CPu were negative at 20Hz stimulation in the Chronos group (**Figure 3C**) and at any rate in the ChR2 group (**Supplementary Figure S5**), while showing positive responses during 75-130Hz in Chronos group (**Figure 3C**). In the ChR2 group, similar analysis revealed two brain regions, the contralateral LGN and SC, that were significantly modulated by pulse repetition rate (**Figure 3D**). Although these regions were not identified as significant in the Chronos group, the extracted time course data revealed similar pulse repetition rate-dependence modulation in these regions within the Chronos group as well the BOLD responses in these two brain regions in the Chronos rats are shown in **Supplementary Figure S6**. In these clusters, we observed that the BOLD signals were higher during 5Hz stimulation compared to the range of 20-130Hz stimulation (**Figure 3E and Supplementary Figure S6**). This pattern suggests that the responses were light-induced visual responses, as indicated in previous research ^26–31^. Therefore, these responses are likely non-therapeutic side effects of optogenetics. In addition, we assessed the voxel-level correlations between BOLD signals and pulse rate-dependent circling behavior effects (**Supplementary Figure S7**) and found a general agreement with these cluster-level results.

### Pulse repetition rate modulated activity in GP and CPu, but not in SNr, were related to the therapeutic effect of DBS

We further determined how STN optogenetic DBS manipulates pathological circling behavior through activity changes in the three identified brain clusters (CPu, GP, and SNr, see **Figure 3B**) using a mediation model with linear mixed-effect regression^32,33^. First, the data demonstrated that the increased pulse repetition rate of STN optogenetic DBS causally manipulated the reduction of normalized angular velocity shown earlier in **Figure 2E** (coefficient = - 0.735, 95% CI = −0.744 – −0.726, P < 0.01; paths c in **Figure 4**). Next, in a multiple mediation model, we set the pulse repetition rate as the independent variable, normalized angular velocity as the dependent variable, and the evoked BOLD fMRI responses in ipsilateral CPu, GP, and SNr as mediators. This analysis revealed that CPu and GP has significant mediation effects (paths a1, b1, a2, b2, all P < 0.01 in **Figure 4**) on the relationship between pulse repetition rate and normalized angular velocity. Critically, such an effect was absent when examining SNr (path b3, n.s. in **Figure 4**). Further, determining the significance of the mediation effect with the Sobel test, suggests both CPu and GP are significant mediators (Sobel z = −14.99, P < 0.01 and −21.82, P < 0.01, respectively). Finally, with CPu and GP incorporated as mediators, the relationship between pulse repetition rate and normalized angular velocity remains significant (path c’, P < 0.01 in **Figure 4**), suggesting CPu and GP are significant partial mediators for the therapeutic effect of STN optogenetic DBS.

**Figure 4.**
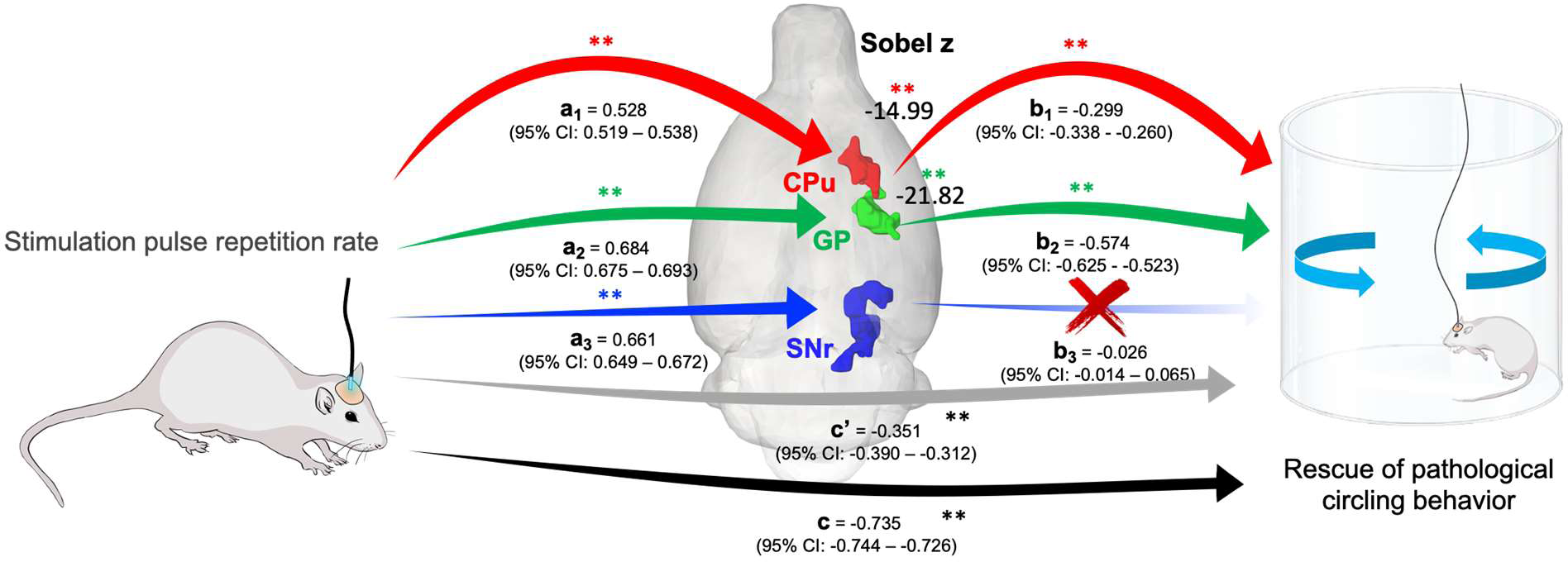
Pulse repetition rate tuning of pathological behavior amelioration is significantly mediated by optogenetic DBS-induced changes in activity in CPu and GP, but not in SNr. Mediation analysis was performed with direct effects (a1, a2, a3, b1, b2, b3, and c) and with mediators (c’). For each significant path, the Sobel test was performed to evaluate the significance of the mediation effect (Sobel z = −14.99 in the path through CPu and Sobel z = −21.82 in the path through GP, p < 0.01). * p < 0.05, ** p < 0.01, and *** p < 0.001; n.s., not significant.

## DISCUSSION

We combined optogenetic DBS and fMRI to map the brain regions specifically modulated by STN feed-forward activity in hemi-parkinsonian rats. We first replicated the behavioral finding in a prior study ^14^ that high pulse repetition rate STN optogenetic DBS using Chronos can ameliorate pathological behavior in hemi-parkinsonian rats. In the same subject groups, we then identified a range of subcortical brain regions, including CPu, GP, and SNr, that were activated by pulse repetition rate-dependent STN optogenetic DBS in rats expressing the fast kinetic Chronos opsin. Notably, pulse rate-dependent activation in these areas was absent (in CPu and GP) or moderate (in SNr) in rats expressing the kinetically slow ChR2 opsin. A mediation model revealed that among the STN feedforward BOLD activity changes found in CPu, GP, and SNr, only CPu and GP exhibit significant partial mediation effects to pulse repetition rate induced amelioration of pathological circling, highlighting the critical roles of the CPu and GP in mediating symptom alleviation by STN DBS. We discuss how our findings may contribute the understanding of mechanisms of DBS in PD.

### Pulse repetition rate dependent BOLD signal changes during STN optogenetics DBS

We used optogenetics to confine the effects of DBS to local STN cell body activation ^14,34^ and leveraged brain-wide fMRI to map the modulated territories ^26,30,35–48^. Optogenetic stimulation modulated the BOLD signal in three major downstream brain areas: SNr, GP and CPu. Notably, the pulse repetition rate dependent effects in GP, SNr, and CPu were only observed in the Chronos group, but not in ChR2 group. The kinetics of ChR2 limit maximum firing rates to ∼40 Hz ^24^, as the slow recovery kinetics of ChR2 can cause depolarization block. In contrast, the frequency-dependent spiking fidelity mediated by fast opsins like Chronos are much closer to those evoked by electrical stimulation ^13,49^. The results in the ChR2 group can thus serve as a positive control for the Chronos effects in activating the STN neurons/axons at high frequencies (>100Hz). In the both groups, we observed light-related stimulus side effects of optogenetic DBS in LGN and SC that were most robust at non-therapeutic, low pulse repetition rates. Activation in LGN and SC are broadly reported in literature studying visual-evoked fMRI responses at low stimulus frequencies as observed in our data ^26–31^.

The principal neurons in STN are glutamatergic ^50–52^, which contrasts with the GABAergic principal neurons found in the rest of the basal ganglia nuclei ^53^. STN receives inhibitory inputs from GP (GP of rodent is homologous to the primate external segment of globus pallidus, GPe) and excitatory inputs from the frontal cortex ^54–56^. STN efferents target intensively the GPe, internal segment of globus pallidus (GPi), and SNr, and sparsely on CPu ^51^. In the current study, we used optogenetics to stimulate the cell bodies in STN, without antidromically activating the cortex or GP (or white matter tracks near the stimulation site). Our findings extend the existing knowledge of anatomical connectivity by revealing robust, pulse repetition rate-dependent functional BOLD responses in SNr, GP, and CPu. The overall rate dependence feature is consistent with previous electrophysiological studies, which found high pulse repletion rate electrical DBS in STN activated efferents to GP, SNr and CPu, and modulated the firing rate and firing pattern in these downstream brain regions ^22,57–67^.

### Pulse repetition rate modulated activity in GP and CPu, but not in SNr, were related to the therapeutic effect of DBS

We employed a mediation model to examine how activity changes in specific brain regions contribute to the improvement of pathological circling behavior. While the BOLD signals in SNr, GP and CPu were all significantly modulated by stimulation pulse repetition rate, the mediation analysis revealed that only GP and CPu, but not SNr, contributed to the reduction of pathological circling.

GP is reciprocally connected with STN, and the GP-STN network generates oscillatory activity that regulates the output of the basal ganglia ^68,69^. In PD, the GP-STN network is altered, leading to pathological beta oscillations (12-30Hz) ^70,71^ that accompany motor symptoms in PD patients ^72^, and which are suppressed by therapeutic STN DBS ^73^. Critically, optogenetic or chemogenetic activation of GP alone was sufficient to attenuate pathological oscillations and/or ameliorate motor dysfunction in PD models ^74–76^. Our result highlights that the change of GP activity plays a critical and causative role in the therapeutic effect of STN DBS. It calls for further studies of the changes of the GP-STN network in understanding the mechanisms of STN DBS for PD.

Our mediation analysis also shows that CPu was involved in the resolution of pathological circling by high pulse rate STN DBS. The modulation of CPu activity by STN DBS may be via direct activation through the sparse projection from STN to CPu, or via indirect activation through pathways inside or outside the basal ganglia. Indeed, a previous study showed that high frequency STN DBS suppressed pathological oscillatory activity in CPu in a PD model ^77^. In addition to directly modulating the CPu activity, STN DBS may also exert its influence by regulating residual dopamine release from substantia nigra pars compacta via the basal ganglia. This notion was supported by experiments showing increased striatal dopamine levels by STN high frequency stimulation in parkinsonian animals ^77–79^.

In contrast to GP and CPu, SNr was not significantly involved in the pulse repetition rate dependent reduction of pathological circling in the present study. Clinically, SNr is not a preferred site (as compared to STN or GPi) for DBS procedures to control PD motor symptoms. One previous study showed that SNr DBS was less effective than STN DBS in improving most of the PD motor symptoms, including akinesia, rigidity, and tremor ^80^. Another study chemically modulated activity in either SNr, GPi or STN in a parkinsonian monkey model, and compared the resulting behavioral effects. It was found that altering the activity of SNr was less effective and less consistent in reducing parkinsonian motor signs. Furthermore, the symptom relieving effects were observed only when manipulating the anterolateral SNr that projects to the ‘motor’ thalamocortical circuit, but not medial SNr which is related to the ‘non-motor’ functions. ^81,82^. The current study selectively stimulated STN neurons, which led to both pulse repetition rate-dependent symptom relief and rate-dependent activation of down-stream regions, including CPu, GP, and SNr, yet the induced SNr activation was not causally related to the improvement of pathological circling. Our results provide evidence that the therapeutic effects of DBS on PD symptoms vary across output nodes of the basal ganglia. The effects on motor symptoms like tremor and akinesia may be via the output of GP. Thus, future research on SNr DBS may focus on its effect on non-motor symptoms of PD ^83,84^ or on gait-related symptoms ^85–90^.

## Conclusion

Numerous studies have underscored the significance of the hyperdirect pathway in therapeutic STN DBS, while downplaying the contribution of STN feed-forward circuits. In a departure from previous approaches that recruited antidromic activation of the hyperdirect pathway or failed to appropriately stimulate STN at high pulse repetition rates, we employed an unbiased, brain-wide fMRI approach coupled with optogenetics using Chronos to identify critical subcortical structures responsible for ameliorating pathological circling behavior in a rat model of PD. Contrary to some prevailing beliefs that subcortical regions downstream of the STN do not play a critical role in STN-DBS therapy, our findings challenged this notion. Specifically, we discovered the causal involvement of two nuclei within the basal ganglia, the GP and the CPu, in mediating the therapeutic effects of STN stimulation. These results highlight the importance of understanding how the reshaping of network activity within the basal ganglia, and the subsequent impact on subcortical basal ganglia output, contribute to the effectiveness of STN DBS therapy. This novel perspective urges further exploration of the STN-GP and STN-CPu pathways, potentially leading to enhanced outcomes in STN DBS interventions for individuals with PD. The interdisciplinary approach presented herein, encompassing optogenetics, fMRI, quantitative behavioral tests, and mediation analysis, serves as a model for investigating diverse neural pathways involved in DBS therapies for a spectrum of neurological and psychiatric disorders.

## ACKNOWLEDGMENT

Funding was provided by the following: National Institutes of Health grant R37NS040894 for WMG; and National Institutes of Health grant R01NS091236 for YIS.

## Author Contributions

Conceptualization: YL, SL, CY, YIS, WMG; Methodology: YL, SL, CY; Investigation: YL, SL, TWW, KD, HK; Formal Analysis & Visualization: YL, SL, LH; original draft: YL, SL, LH, YIS, WMG; review and final editing: YL, SL, YIS, WMG.

## EXPERIMENTAL PROCEDURES

### Animals and surgery

All animal care and experimental procedures were approved by the University of North Carolina Institutional Animal Care and Use Committee. Fourteen female Sprague Dawley rats (weight 250−275g, ∼90 days old) were used in this study. The animals were housed in a temperature-controlled environment on a 12-h light-dark cycle and provided with ad-libitum food and water. Rats received either rAAV5-CAMKII-Chronos-GFP (n=8) or AAV-CAMKII-ChR2-GFP (n=6) injection. There was an unexpected failure of the MRI system during the research, and data collection in five rats were abandoned due to the concerns of headcap integrity and health condition because of the long delay. The remaining nine rats (Chronos group n = 5, ChR2 group n = 4) were used both in methamphetamine-induced circling measurements and functional imaging experiments.

Surgery was conducted under anesthesia with sevoflurane (induced at 7% and maintained at 3.25%). The animals were fixed in a stereotaxic frame. The body temperature was maintained at 37°C by a heated water blanket. Meloxicam (2.0 mg/kg, s.q.) and Bupivacaine (0.2ml of 0.25%, s.q.) were administered for analgesia. Dexamethasone (0.5 mg/kg, s.q.) was administered to reduce cerebral edema. Craniotomies were performed over the STN (3.6 mm posterior and 2.6 mm lateral from bregma) and medial forebrain bundle (MFB, 2.0 mm posterior and 2.0 mm lateral from bregma) in the left hemisphere. rAAV5-CAMKII-Chronos-GFP (1.8 × 10^13^ vg/ml) or AAV-CAMKII-ChR2-GFP (4.6 × 10^12^ vg/ml, University of North Carolina vector core) was injected into the STN (depth ∼7.0 mm from cortical surface; 0.6 uL at a rate of 0.15 uL/min). Six µL of 6-hydroxydopamine (6-OHDA) (2.5 mg/ml dissolved in 0.2% ascorbic acid in saline) was injected into the MFB (depth 7.5 mm from cortical surface) at a rate of 1.5 uL/min to induce unilateral dopaminergic lesion of the substantia nigra pars compacta (SNc). Desipramine (5 mg/kg) and pargyline (50 mg/kg) were administered intraperitoneally thirty minutes before the 6-OHDA injection to protect noradrenergic neurons and inhibit monoamine oxidase, respectively. An optical fiber (200 µm core diameter, 0.37 NA, Precision Fiber Products) was implanted into the STN 300µm dorsal to the virus injection site. The optical fiber was then fixed to the skull by MRI compatible bone screws and dental cement.

### Behavioral test and optogenetic stimulation

The success of the dopaminergic lesion and effects of DBS were assessed using methamphetamine-induced circling ^91^ starting four weeks after the surgery. The animals were administered 1.875 mg/kg of methamphetamine (intraperitoneal, 1.25 mg/ml dissolved in saline) and then placed in a dark cylindrical chamber 30 cm in diameter. The locomotor activity of the rat was captured by an infrared camera. Six different conditions were tested in random order: no optical stimulus, 5Hz, 20Hz, 75Hz, 100Hz or 130Hz optical stimuli. In each test condition, laser light (473-nm, Shanghai Laser) was applied via an optical fiber. The laser power was 10 mW (calibrated as output power at the end of the fiber) and the pulse width was 1 ms. The duration of stimulation was 10 seconds with 20 s intervals between epochs of stimulation. Each stimulation condition was repeated six times.

### fMRI Data collection

The fMRI scans were conducted one week after the circling tests following the procedures established in several recent studies ^15,92–94^. Briefly, rats were intubated and ventilated under dexmedetomidine (0.025mg/kg/hour, i.p infusion) and 0.5% isoflurane. Pancuronium bromide (0.5 mg/kg/h, i.p infusion) was administered for paralysis. Heart rate and oxygen saturation level were continuously monitored by a MouseOX Plus system (STARR Life Science Corp.) and maintained at 350-400 bpm and 95%, respectively. Body temperature was maintained at 37°C by a circulating water pad. End-tidal CO_2_ was monitored by a capnometer (Surgivet, Smith Medical) and stabilized within 2.6 - 3.2%. MR images were collected through the UNC Center for Animal MRI (CAMRI) service on a Bruker BioSpec 9.4-Tesla, 30 cm bore system (Bruker BioSpin Corp., Billerica, MA) with Paravision 6.0.1 on an AVANCE II console and house-built single-loop coil. Magnetic field homogeneity was first optimized by global shimming, followed by local second-order shims using a MAPSHIM protocol. For each subject, the BOLD fMRI data were acquired using a 2D multi-slice, single-shot, gradient echo-planar imaging (EPI) sequence: repetition time (TR) = 2000 ms, echo time (TE) = 14 ms, bandwidth = 250 kHz, flip angle = 70 degrees, field of view (FOV) = 28.8 x 28.8 mm, matrix size = 72 x 72, slice number = 32, and slice 130Hz) were delivered in random order. Three repeated scan sessions of 20 s rest period followed by six repetitions of the optical stimulation train, including 10 s of stimulation and 20 s rest period block, were applied for each pulse repetition rate.

### Data analysis

The position of the nose and the base of tail over time were extracted from the video recording of the methamphetamine-induced circling using TopScan analysis software (Clever Systems). The normalized angular velocity and linear speed of the body for each trial were calculated by dividing the angular velocity or linear speed during the 10 s stimulation-on period by the averaged angular velocities or linear speeds during the 20 s pre-stimulation and post-stimulation periods immediately before and after the stimulation-on period. Normalized angular velocity or linear speed for each pulse repetition rate were then computed by averaging across all repetitions.

For MRI data analysis, all EPI data were first corrected for slice timing, followed by motion correction using AFNI ^95^. Each 4D dataset was then averaged across time to improve signal-to-noise ratios (SNR) for subsequent skull stripping. The brain mask for each time-averaged EPI image was manually drawn using ITK-snap ^96^. The skull-stripped data were then spatially normalized to the T2 anatomical rat brain template using ANTs SyN diffeomorphic registration ^97^ before further group-level analysis. General Linear Model (GLM) analysis was conducted using the 3dDeconvolve tool in AFNI to generate evoked response maps corresponding to the optogenetic stimulation. AFNI’s statistical package was used to conduct group-level statistics. First, the effects of stimulation frequency, opsins, and the interaction between stimulation frequency and opsins were evaluated with a linear mixed model using 3dLME. The intercept and repeated scan trials were set as random effects. We also designed a contrast matrix and applied it to the estimated linear mixed model for evaluating the effect size of each factor and testing the significant differences between their averages. Specifically, we applied an ANOVA to test for significant differences between the averages of each factor, including the main effects of stimulation frequency, opsin, and their interaction. For the multiple comparison correction, the spatial autocorrelation function (sACF) was estimated by fitting the mixed model of gaussian and mono-exponential function for the residual of the linear mixed model analysis (using 3dFWHMx). Subsequently, the probability of false positive clusters was estimated from the sACF to identify the minimal size of significant clusters at 5% of the false positive rate (using 3dClustSim). The clusters larger than the estimated cluster size were determined as significant findings.

To explore how optogenetics STN DBS affects pathological circling behavior through brain activations, a mediation model using linear mixed-effect regression (LMER)^98^ was constructed. The study first calculated the direct relationships between pulse repetition rate and normalized angular velocity, with the LMER model used to estimate the coefficient between the two variables and handle longitudinal data. Random intercept and subject effects were included in each LMER model to account for temporal correlation. Then, the mediation model estimated the indirect relationship between pulse repetition rate and normalized angular velocity, with three identified DBS stimulated-related brain activations as mediations. In the mediation model, pulse repetition rate served as the independent variable, normalized angular velocity as the dependent variable, and the three coefficients of BOLD response from SNr, GP and CPu as mediator variables. We also employed 1,000 bootstrapping tests to assess the significance of each pathway, and the Sobel test was used to evaluate the mediation effect of each significant path ^99^.

### Histology

At the conclusion of the MRI scan, rats were deeply anesthetized with phenytoin sodium (Euthasol) and transcardially perfused with saline followed by 4% paraformaldehyde. The brain was removed and stored overnight in 4% paraformaldehyde and transferred to a 10% sucrose (in DI water) for 4 h and then 30% sucrose solution for 2–3 days, until brains sunk. Brains were cut at 40 μm thick sections on a freezing microtome horizontally and mounted on glass slides for fluorescent imaging. Tyrosine hydroxylase (TH) immunohistochemistry was used to evaluate the degree of degeneration of dopaminergic neurons in the SNc. Briefly, brain slices were rinsed and incubated in PBST (0.3% of Triton-X in PBS) with 3% goat serum for 30 minutes at room temperature. The sections were then incubated in anti-TH antibody (1:1000, AB152, Abcam, CA, USA) at 4 °C for 16 hours. After rinsing in PBST 3 times for 10 minutes each time, the sections were incubated in secondary antibody (1:1000, goat anti-rabbit Alexa Fluor 546) for 2 hours in room temperature before being rinsed and mounted to slides. Vectashield mounting medium with DAPI stain (Vector laboratories, Item # H−1200) was used to provide a cell body counterstain. To confirm the expression of Chronos-GFP, brain sections were directly mounted. To validate the cell-type specific expression of Chronos-GFP, brain sections were stained for CaMKII. Slices were incubated with primary anti-CaMKII antibody (1:300, Millipore #05-532) with 10% goat serum and 0.25% Triton X−100 overnight at 4 °C. Second antibodies (Alexa Fluor 594 goat anti-mouse IgGI) were used to visualize CaMKII. All sections were mounted with DAPI-FluoroMount-G (Southern Biotech) and were imaged with a 20x oil lens with Olympus FV3000RS confocal microscope or Nikon Eclipse Ti2 microscope. The locations of the optical fibers were determined by registering the MR anatomical images to a rat brain atlas ^100^.

### Statistics

A two-way ANOVA analysis was used to test the significance of pulse repetition rate effect and injected virus type on angular velocity or linear speed in the methamphetamine-induced circling test. *Post hoc* paired comparisons with Tukey-Kramer method were used to compare the difference between two different pulse repetition rate conditions. All statistic results are represented as mean ± standard deviation unless otherwise indicated.

## Data Availability

The datasets generated during and/or analyzed during the current study are available from the corresponding author W.M.G. upon request.

## REFERENCES

1. Schuepbach, W.M., Rau, J., Knudsen, K., Volkmann, J., Krack, P., Timmermann, L., Halbig, T.D., Hesekamp, H., Navarro, S.M., Meier, N., et al. (2013). Neurostimulation for Parkinson’s disease with early motor complications. N Engl J Med 368, 610–622. 10.1056/NEJMoa1205158.

2. Hacker, M.L., DeLong, M.R., Turchan, M., Heusinkveld, L.E., Ostrem, J.L., Molinari, A.L., Currie, A.D., Konrad, P.E., Davis, T.L., Phibbs, F.T., et al. (2018). Effects of deep brain stimulation on rest tremor progression in early stage Parkinson disease. Neurology 91, e463–e471. 10.1212/WNL.0000000000005903.

3. Hacker, M.L., Turchan, M., Heusinkveld, L.E., Currie, A.D., Millan, S.H., Molinari, A.L., Konrad, P.E., Davis, T.L., Phibbs, F.T., Hedera, P., et al. (2020). Deep brain stimulation in early-stage Parkinson disease: Five-year outcomes. Neurology 95, e393–e401. 10.1212/WNL.0000000000009946.

4. Asanuma, K., Tang, C., Ma, Y., Dhawan, V., Mattis, P., Edwards, C., Kaplitt, M.G., Feigin, A., and Eidelberg, D. (2006). Network modulation in the treatment of Parkinson’s disease. Brain 129, 2667–2678. 10.1093/brain/awl162.

5. Grafton, S.T., Turner, R.S., Desmurget, M., Bakay, R., Delong, M., Vitek, J., and Crutcher, M. (2006). Normalizing motor-related brain activity: subthalamic nucleus stimulation in Parkinson disease. Neurology 66, 1192–1199. 10.1212/01.wnl.0000214237.58321.c3.

6. Kahan, J., Mancini, L., Urner, M., Friston, K., Hariz, M., Holl, E., White, M., Ruge, D., Jahanshahi, M., Boertien, T., et al. (2012). Therapeutic subthalamic nucleus deep brain stimulation reverses cortico-thalamic coupling during voluntary movements in Parkinson’s disease. PLoS One 7, e50270. 10.1371/journal.pone.0050270.

7. Phillips, M.D., Baker, K.B., Lowe, M.J., Tkach, J.A., Cooper, S.E., Kopell, B.H., and Rezai, A.R. (2006). Parkinson disease: pattern of functional MR imaging activation during deep brain stimulation of subthalamic nucleus--initial experience. Radiology 239, 209–216. 10.1148/radiol.2391041990.

8. Stefurak, T., Mikulis, D., Mayberg, H., Lang, A.E., Hevenor, S., Pahapill, P., Saint-Cyr, J., and Lozano, A. (2003). Deep brain stimulation for Parkinson’s disease dissociates mood and motor circuits: a functional MRI case study. Mov Disord 18, 1508–1516. 10.1002/mds.10593.

9. Kim, C.K., Adhikari, A., and Deisseroth, K. (2017). Integration of optogenetics with complementary methodologies in systems neuroscience. Nat Rev Neurosci 18, 222–235. 10.1038/nrn.2017.15.

10. Gradinaru, V., Mogri, M., Thompson, K.R., Henderson, J.M., and Deisseroth, K. (2009). Optical deconstruction of parkinsonian neural circuitry. Science 324, 354–359. 10.1126/science.1167093.

11. Sanders, T.H., and Jaeger, D. (2016). Optogenetic stimulation of cortico-subthalamic projections is sufficient to ameliorate bradykinesia in 6-ohda lesioned mice. Neurobiol Dis 95, 225–237. 10.1016/j.nbd.2016.07.021.

12. Boyden, E.S., Zhang, F., Bamberg, E., Nagel, G., and Deisseroth, K. (2005). Millisecond-timescale, genetically targeted optical control of neural activity. Nat Neurosci 8, 1263–1268. 10.1038/nn1525.

13. Hight, A.E., Kozin, E.D., Darrow, K., Lehmann, A., Boyden, E., Brown, M.C., and Lee, D.J. (2015). Superior temporal resolution of Chronos versus channelrhodopsin−2 in an optogenetic model of the auditory brainstem implant. Hear Res 322, 235–241. 10.1016/j.heares.2015.01.004.

14. Yu, C., Cassar, I.R., Sambangi, J., and Grill, W.M. (2020). Frequency-Specific Optogenetic Deep Brain Stimulation of Subthalamic Nucleus Improves Parkinsonian Motor Behaviors. J Neurosci 40, 4323–4334. 10.1523/JNEUROSCI.3071-19.2020.

15. Cerri, D.H., Albaugh, D.L., Walton, L.R., Katz, B., Wang, T.-W., Chao, T.-H.H., Zhang, W., Nonneman, R.J., Jiang, J., and Lee, S.-H. (2023). Distinct neurochemical influences on fMRI response polarity in the striatum. BioRxiv, 2023.2002. 2020.529283.

16. Lai, H.Y., Younce, J.R., Albaugh, D.L., Kao, Y.C., and Shih, Y.Y. (2014). Functional MRI reveals frequency-dependent responses during deep brain stimulation at the subthalamic nucleus or internal globus pallidus. Neuroimage 84, 11–18. 10.1016/j.neuroimage.2013.08.026.

17. Lee, J.H., Liu, Q., and Dadgar-Kiani, E. (2022). Solving brain circuit function and dysfunction with computational modeling and optogenetic fMRI. Science 378, 493–499. 10.1126/science.abq3868.

18. Lee, J.Y., You, T., Woo, C.W., and Kim, S.G. (2022). Optogenetic fMRI for Brain-Wide Circuit Analysis of Sensory Processing. Int J Mol Sci 23. 10.3390/ijms232012268.

19. Menon, V., Cerri, D., Lee, B., Yuan, R., Lee, S.H., and Shih, Y.I. (2023). Optogenetic stimulation of anterior insular cortex neurons in male rats reveals causal mechanisms underlying suppression of the default mode network by the salience network. Nat Commun 14, 866. 10.1038/s41467-023-36616-8.

20. Van Den Berge, N., Albaugh, D.L., Salzwedel, A., Vanhove, C., Van Holen, R., Gao, W., Stuber, G.D., and Shih, Y.I. (2017). Functional circuit mapping of striatal output nuclei using simultaneous deep brain stimulation and fMRI. Neuroimage 146, 1050–1061. 10.1016/j.neuroimage.2016.10.049.

21. Zhao, S., Li, G., Tong, C., Chen, W., Wang, P., Dai, J., Fu, X., Xu, Z., Liu, X., Lu, L., et al. (2020). Full activation pattern mapping by simultaneous deep brain stimulation and fMRI with graphene fiber electrodes. Nat Commun 11, 1788. 10.1038/s41467-020-15570-9.

22. McConnell, G.C., So, R.Q., Hilliard, J.D., Lopomo, P., and Grill, W.M. (2012). Effective deep brain stimulation suppresses low-frequency network oscillations in the basal ganglia by regularizing neural firing patterns. J Neurosci 32, 15657–15668. 10.1523/JNEUROSCI.2824-12.2012.

23. Moro, E., Esselink, R.J., Xie, J., Hommel, M., Benabid, A.L., and Pollak, P. (2002). The impact on Parkinson’s disease of electrical parameter settings in STN stimulation. Neurology 59, 706–713. 10.1212/wnl.59.5.706.

24. Klapoetke, N.C., Murata, Y., Kim, S.S., Pulver, S.R., Birdsey-Benson, A., Cho, Y.K., Morimoto, T.K., Chuong, A.S., Carpenter, E.J., Tian, Z., et al. (2014). Independent optical excitation of distinct neural populations. Nat Methods 11, 338–346. 10.1038/nmeth.2836.

25. So, R.Q., McConnell, G.C., and Grill, W.M. (2017). Frequency-dependent, transient effects of subthalamic nucleus deep brain stimulation on methamphetamine-induced circling and neuronal activity in the hemiparkinsonian rat. Behav Brain Res 320, 119–127. 10.1016/j.bbr.2016.12.003.

26. Decot, H.K., Namboodiri, V.M., Gao, W., McHenry, J.A., Jennings, J.H., Lee, S.H., Kantak, P.A., Jill Kao, Y.C., Das, M., Witten, I.B., et al. (2017). Coordination of Brain-Wide Activity Dynamics by Dopaminergic Neurons. Neuropsychopharmacology 42, 615–627. 10.1038/npp.2016.151.

27. Dinh, T.N.A., Jung, W.B., Shim, H.J., and Kim, S.G. (2021). Characteristics of fMRI responses to visual stimulation in anesthetized vs. awake mice. Neuroimage 226, 117542. 10.1016/j.neuroimage.2020.117542.

28. Gil, R., Fernandes, F.F., and Shemesh, N. (2021). Neuroplasticity-driven timing modulations revealed by ultrafast functional magnetic resonance imaging. Neuroimage 225, 117446. 10.1016/j.neuroimage.2020.117446.

29. Lee, H.L., Li, Z., Coulson, E.J., and Chuang, K.H. (2019). Ultrafast fMRI of the rodent brain using simultaneous multi-slice EPI. Neuroimage 195, 48–58. 10.1016/j.neuroimage.2019.03.045.

30. Leong, A.T.L., Gu, Y., Chan, Y.S., Zheng, H., Dong, C.M., Chan, R.W., Wang, X., Liu, Y., Tan, L.H., and Wu, E.X. (2019). Optogenetic fMRI interrogation of brain-wide central vestibular pathways. Proc Natl Acad Sci U S A 116, 10122–10129. 10.1073/pnas.1812453116.

31. Schmid, F., Wachsmuth, L., Albers, F., Schwalm, M., Stroh, A., and Faber, C. (2017). True and apparent optogenetic BOLD fMRI signals. Magn Reson Med 77, 126–136. 10.1002/mrm.26095.

32. Chén, O.Y., Crainiceanu, C., Ogburn, E.L., Caffo, B.S., Wager, T.D., and Lindquist, M.A. (2018). High-dimensional multivariate mediation with application to neuroimaging data. Biostatistics 19, 121–136.

33. Oyarzabal, E.A., Hsu, L.-M., Das, M., Chao, T.-H.H., Zhou, J., Song, S., Zhang, W., Smith, K.G., Sciolino, N.R., and Evsyukova, I.Y. (2022). Chemogenetic stimulation of tonic locus coeruleus activity strengthens the default mode network. Science advances 8, eabm9898.

34. Zhang, F., Gradinaru, V., Adamantidis, A.R., Durand, R., Airan, R.D., de Lecea, L., and Deisseroth, K. (2010). Optogenetic interrogation of neural circuits: technology for probing mammalian brain structures. Nat Protoc 5, 439–456. 10.1038/nprot.2009.226.

35. Chen, Y., Pais-Roldan, P., Chen, X., Frosz, M.H., and Yu, X. (2019). MRI-guided robotic arm drives optogenetic fMRI with concurrent Ca(2+) recording. Nat Commun 10, 2536. 10.1038/s41467-019-10450-3.

36. Ferenczi, E.A., Zalocusky, K.A., Liston, C., Grosenick, L., Warden, M.R., Amatya, D., Katovich, K., Mehta, H., Patenaude, B., Ramakrishnan, C., et al. (2016). Prefrontal cortical regulation of brainwide circuit dynamics and reward-related behavior. Science 351, aac9698. 10.1126/science.aac9698.

37. Gozzi, A., and Zerbi, V. (2023). Modeling Brain Dysconnectivity in Rodents. Biol Psychiatry 93, 419–429. 10.1016/j.biopsych.2022.09.008.

38. Grandjean, J., Corcoba, A., Kahn, M.C., Upton, A.L., Deneris, E.S., Seifritz, E., Helmchen, F., Mann, E.O., Rudin, M., and Saab, B.J. (2019). A brain-wide functional map of the serotonergic responses to acute stress and fluoxetine. Nat Commun 10, 350. 10.1038/s41467-018-08256-w.

39. Grimm, C., Frassle, S., Steger, C., von Ziegler, L., Sturman, O., Shemesh, N., Peleg-Raibstein, D., Burdakov, D., Bohacek, J., Stephan, K.E., et al. (2021). Optogenetic activation of striatal D1R and D2R cells differentially engages downstream connected areas beyond the basal ganglia. Cell Rep 37, 110161. 10.1016/j.celrep.2021.110161.

40. Jung, W.B., Jiang, H., Lee, S., and Kim, S.G. (2022). Dissection of brain-wide resting-state and functional somatosensory circuits by fMRI with optogenetic silencing. Proc Natl Acad Sci U S A 119. 10.1073/pnas.2113313119.

41. Kim, S., Moon, H.S., Vo, T.T., Kim, C.H., Im, G.H., Lee, S., Choi, M., and Kim, S.G. (2023). Whole-brain mapping of effective connectivity by fMRI with cortex-wide patterned optogenetics. Neuron 111, 1732–1747 e1736. 10.1016/j.neuron.2023.03.002.

42. Lee, J.H., Durand, R., Gradinaru, V., Zhang, F., Goshen, I., Kim, D.S., Fenno, L.E., Ramakrishnan, C., and Deisseroth, K. (2010). Global and local fMRI signals driven by neurons defined optogenetically by type and wiring. Nature 465, 788–792. 10.1038/nature09108.

43. Lee, J.Y., You, T., Lee, C.H., Im, G.H., Seo, H., Woo, C.W., and Kim, S.G. (2022). Role of anterior cingulate cortex inputs to periaqueductal gray for pain avoidance. Curr Biol 32, 2834–2847 e2835. 10.1016/j.cub.2022.04.090.

44. Liang, Z., Watson, G.D., Alloway, K.D., Lee, G., Neuberger, T., and Zhang, N. (2015). Mapping the functional network of medial prefrontal cortex by combining optogenetics and fMRI in awake rats. Neuroimage 117, 114–123. 10.1016/j.neuroimage.2015.05.036.

45. Mandino, F., Vrooman, R.M., Foo, H.E., Yeow, L.Y., Bolton, T.A.W., Salvan, P., Teoh, C.L., Lee, C.Y., Beauchamp, A., Luo, S., et al. (2022). A triple-network organization for the mouse brain. Mol Psychiatry 27, 865–872. 10.1038/s41380-021-01298-5.

46. Ryali, S., Shih, Y.Y., Chen, T., Kochalka, J., Albaugh, D., Fang, Z., Supekar, K., Lee, J.H., and Menon, V. (2016). Combining optogenetic stimulation and fMRI to validate a multivariate dynamical systems model for estimating causal brain interactions. Neuroimage 132, 398–405. 10.1016/j.neuroimage.2016.02.067.

47. Toi, P.T., Jang, H.J., Min, K., Kim, S.P., Lee, S.K., Lee, J., Kwag, J., and Park, J.Y. (2022). In vivo direct imaging of neuronal activity at high temporospatial resolution. Science 378, 160–168. 10.1126/science.abh4340.

48. Cover, C.G., Kesner, A.J., Ukani, S., Stein, E.A., Ikemoto, S., Yang, Y., and Lu, H. (2021). Whole brain dynamics during optogenetic self-stimulation of the medial prefrontal cortex in mice. Commun Biol 4, 66. 10.1038/s42003-020-01612-x.

49. Mattis, J., Tye, K.M., Ferenczi, E.A., Ramakrishnan, C., O’Shea, D.J., Prakash, R., Gunaydin, L.A., Hyun, M., Fenno, L.E., Gradinaru, V., et al. (2011). Principles for applying optogenetic tools derived from direct comparative analysis of microbial opsins. Nat Methods 9, 159–172. 10.1038/nmeth.1808.

50. Albin, R.L., Aldridge, J.W., Young, A.B., and Gilman, S. (1989). Feline subthalamic nucleus neurons contain glutamate-like but not GABA-like or glycine-like immunoreactivity. Brain Res 491, 185–188. 10.1016/0006-8993(89)90103-0.

51. Kita, H., and Kitai, S.T. (1987). Efferent projections of the subthalamic nucleus in the rat: light and electron microscopic analysis with the PHA-L method. J Comp Neurol 260, 435–452. 10.1002/cne.902600309.

52. Smith, Y., and Parent, A. (1988). Neurons of the subthalamic nucleus in primates display glutamate but not GABA immunoreactivity. Brain Res 453, 353–356. 10.1016/0006-8993(88)90177-1.

53. Lanciego, J.L., Luquin, N., and Obeso, J.A. (2012). Functional neuroanatomy of the basal ganglia. Cold Spring Harb Perspect Med 2, a009621. 10.1101/cshperspect.a009621.

54. Kita, H., Chang, H.T., and Kitai, S.T. (1983). The morphology of intracellularly labeled rat subthalamic neurons: a light microscopic analysis. J Comp Neurol 215, 245–257. 10.1002/cne.902150302.

55. Nakanishi, H., Kita, H., and Kitai, S.T. (1987). Electrical membrane properties of rat subthalamic neurons in an in vitro slice preparation. Brain Res 437, 35–44. 10.1016/0006-8993(87)91524-1.

56. Nakanishi, H., Kita, H., and Kitai, S.T. (1988). An N-methyl-D-aspartate receptor mediated excitatory postsynaptic potential evoked in subthalamic neurons in an in vitro slice preparation of the rat. Neurosci Lett 95, 130–136. 10.1016/0304-3940(88)90645-3.

57. Adam, E.M., Brown, E.N., Kopell, N., and McCarthy, M.M. (2022). Deep brain stimulation in the subthalamic nucleus for Parkinson’s disease can restore dynamics of striatal networks. Proc Natl Acad Sci U S A 119, e2120808119. 10.1073/pnas.2120808119.

58. Benazzouz, A., Piallat, B., Pollak, P., and Benabid, A.L. (1995). Responses of substantia nigra pars reticulata and globus pallidus complex to high frequency stimulation of the subthalamic nucleus in rats: electrophysiological data. Neurosci Lett 189, 77–80. 10.1016/0304-3940(95)11455-6.

59. Galati, S., Mazzone, P., Fedele, E., Pisani, A., Peppe, A., Pierantozzi, M., Brusa, L., Tropepi, D., Moschella, V., Raiteri, M., et al. (2006). Biochemical and electrophysiological changes of substantia nigra pars reticulata driven by subthalamic stimulation in patients with Parkinson’s disease. Eur J Neurosci 23, 2923–2928. 10.1111/j.1460-9568.2006.04816.x.

60. Hammond, C., Deniau, J.M., Rizk, A., and Feger, J. (1978). Electrophysiological demonstration of an excitatory subthalamonigral pathway in the rat. Brain Res 151, 235–244. 10.1016/0006-8993(78)90881-8.

61. Hashimoto, T., Elder, C.M., Okun, M.S., Patrick, S.K., and Vitek, J.L. (2003). Stimulation of the subthalamic nucleus changes the firing pattern of pallidal neurons. J Neurosci 23, 1916–1923. 10.1523/JNEUROSCI.23-05-01916.2003.

62. Maurice, N., Thierry, A.M., Glowinski, J., and Deniau, J.M. (2003). Spontaneous and evoked activity of substantia nigra pars reticulata neurons during high-frequency stimulation of the subthalamic nucleus. J Neurosci 23, 9929–9936. 10.1523/JNEUROSCI.23-30-09929.2003.

63. Min, H.K., Ross, E.K., Jo, H.J., Cho, S., Settell, M.L., Jeong, J.H., Duffy, P.S., Chang, S.Y., Bennet, K.E., Blaha, C.D., and Lee, K.H. (2016). Dopamine Release in the Nonhuman Primate Caudate and Putamen Depends upon Site of Stimulation in the Subthalamic Nucleus. J Neurosci 36, 6022–6029. 10.1523/JNEUROSCI.0403-16.2016.

64. Reese, R., Leblois, A., Steigerwald, F., Potter-Nerger, M., Herzog, J., Mehdorn, H.M., Deuschl, G., Meissner, W.G., and Volkmann, J. (2011). Subthalamic deep brain stimulation increases pallidal firing rate and regularity. Exp Neurol 229, 517–521. 10.1016/j.expneurol.2011.01.020.

65. Shi, L.H., Luo, F., Woodward, D.J., and Chang, J.Y. (2006). Basal ganglia neural responses during behaviorally effective deep brain stimulation of the subthalamic nucleus in rats performing a treadmill locomotion test. Synapse 59, 445–457. 10.1002/syn.20261.

66. Tai, C.H., Boraud, T., Bezard, E., Bioulac, B., Gross, C., and Benazzouz, A. (2003). Electrophysiological and metabolic evidence that high-frequency stimulation of the subthalamic nucleus bridles neuronal activity in the subthalamic nucleus and the substantia nigra reticulata. FASEB J 17, 1820–1830. 10.1096/fj.03-0163com.

67. Windels, F., Bruet, N., Poupard, A., Urbain, N., Chouvet, G., Feuerstein, C., and Savasta, M. (2000). Effects of high frequency stimulation of subthalamic nucleus on extracellular glutamate and GABA in substantia nigra and globus pallidus in the normal rat. Eur J Neurosci 12, 4141–4146. 10.1046/j.1460-9568.2000.00296.x.

68. Bevan, M.D., Magill, P.J., Terman, D., Bolam, J.P., and Wilson, C.J. (2002). Move to the rhythm: oscillations in the subthalamic nucleus-external globus pallidus network. Trends Neurosci 25, 525–531. 10.1016/s0166-2236(02)02235-x.

69. Plenz, D., and Kital, S.T. (1999). A basal ganglia pacemaker formed by the subthalamic nucleus and external globus pallidus. Nature 400, 677–682. 10.1038/23281.

70. Chu, H.Y., Atherton, J.F., Wokosin, D., Surmeier, D.J., and Bevan, M.D. (2015). Heterosynaptic regulation of external globus pallidus inputs to the subthalamic nucleus by the motor cortex. Neuron 85, 364–376. 10.1016/j.neuron.2014.12.022.

71. Mallet, N., Pogosyan, A., Marton, L.F., Bolam, J.P., Brown, P., and Magill, P.J. (2008). Parkinsonian beta oscillations in the external globus pallidus and their relationship with subthalamic nucleus activity. J Neurosci 28, 14245–14258. 10.1523/JNEUROSCI.4199-08.2008.

72. Neumann, W.J., Degen, K., Schneider, G.H., Brucke, C., Huebl, J., Brown, P., and Kuhn, A.A. (2016). Subthalamic synchronized oscillatory activity correlates with motor impairment in patients with Parkinson’s disease. Mov Disord 31, 1748–1751. 10.1002/mds.26759.

73. Quinn, E.J., Blumenfeld, Z., Velisar, A., Koop, M.M., Shreve, L.A., Trager, M.H., Hill, B.C., Kilbane, C., Henderson, J.M., and Bronte-Stewart, H. (2015). Beta oscillations in freely moving Parkinson’s subjects are attenuated during deep brain stimulation. Mov Disord 30, 1750–1758. 10.1002/mds.26376.

74. Assaf, F., and Schiller, Y. (2019). A chemogenetic approach for treating experimental Parkinson’s disease. Mov Disord 34, 469–479. 10.1002/mds.27554.

75. Crompe, B., Aristieta, A., Leblois, A., Elsherbiny, S., Boraud, T., and Mallet, N.P. (2020). The globus pallidus orchestrates abnormal network dynamics in a model of Parkinsonism. Nat Commun 11, 1570. 10.1038/s41467-020-15352-3.

76. Mastro, K.J., Zitelli, K.T., Willard, A.M., Leblanc, K.H., Kravitz, A.V., and Gittis, A.H. (2017). Cell-specific pallidal intervention induces long-lasting motor recovery in dopamine-depleted mice. Nat Neurosci 20, 815–823. 10.1038/nn.4559.

77. Zhang, Y., Xu, S., Xiao, G., Song, Y., Gao, F., Wang, M., Zhao, H., Xing, G., and Cai, X. (2019). High frequency stimulation of subthalamic nucleus synchronously modulates primary motor cortex and caudate putamen based on dopamine concentration and electrophysiology activities using microelectrode arrays in Parkinson’s disease rats. Sensors and Actuators B: Chemical 301, 127126.

78. Bruet, N., Windels, F., Bertrand, A., Feuerstein, C., Poupard, A., and Savasta, M. (2001). High frequency stimulation of the subthalamic nucleus increases the extracellular contents of striatal dopamine in normal and partially dopaminergic denervated rats. J Neuropathol Exp Neurol 60, 15–24. 10.1093/jnen/60.1.15.

79. Gale, J.T., Lee, K.H., Amirnovin, R., Roberts, D.W., Williams, Z.M., Blaha, C.D., and Eskandar, E.N. (2013). Electrical stimulation-evoked dopamine release in the primate striatum. Stereotact Funct Neurosurg 91, 355–363. 10.1159/000351523.

80. Chastan, N., Westby, G.W., Yelnik, J., Bardinet, E., Do, M.C., Agid, Y., and Welter, M.L. (2009). Effects of nigral stimulation on locomotion and postural stability in patients with Parkinson’s disease. Brain 132, 172–184. 10.1093/brain/awn294.

81. Wichmann, T., Baron, M.S., and DeLong, M.R. (1994). Local inactivation of the sensorimotor territories of the internal segment of the globus pallidus and the subthalamic nucleus alleviates parkinsonian motor signs in MPTP treated monkeys. the basal ganglia IV: New ideas and data on structure and function, 357–363.

82. Wichmann, T., Kliem, M.A., and DeLong, M.R. (2001). Antiparkinsonian and behavioral effects of inactivation of the substantia nigra pars reticulata in hemiparkinsonian primates. Exp Neurol 167, 410–424. 10.1006/exnr.2000.7572.

83. Cascella, N., Butala, A.A., Mills, K., Kim, M.J., Salimpour, Y., Wojtasievicz, T., Hwang, B., Cullen, B., Figee, M., Moran, L., et al. (2021). Deep Brain Stimulation of the Substantia Nigra Pars Reticulata for Treatment-Resistant Schizophrenia: A Case Report. Biol Psychiatry 90, e57–e59. 10.1016/j.biopsych.2021.03.007.

84. Zhang, L., Meng, S., Chen, W., Chen, Y., Huang, E., Zhang, G., Liang, Y., Ding, Z., Xue, Y., Chen, Y., et al. (2021). High-Frequency Deep Brain Stimulation of the Substantia Nigra Pars Reticulata Facilitates Extinction and Prevents Reinstatement of Methamphetamine-Induced Conditioned Place Preference. Front Pharmacol 12, 705813. 10.3389/fphar.2021.705813.

85. Avila, I., Parr-Brownlie, L.C., Brazhnik, E., Castaneda, E., Bergstrom, D.A., and Walters, J.R. (2010). Beta frequency synchronization in basal ganglia output during rest and walk in a hemiparkinsonian rat. Exp Neurol 221, 307–319. 10.1016/j.expneurol.2009.11.016.

86. Burbaud, P., Bonnet, B., Guehl, D., Lagueny, A., and Bioulac, B. (1998). Movement disorders induced by gamma-aminobutyric agonist and antagonist injections into the internal globus pallidus and substantia nigra pars reticulata of the monkey. Brain Res 780, 102–107. 10.1016/s0006-8993(97)01158-x.

87. Golfre Andreasi, N., Rispoli, V., Contaldi, E., Colucci, F., Mongardi, L., Cavallo, M.A., and Sensi, M. (2020). Deep brain stimulation and refractory freezing of gait in Parkinson’s disease: Improvement with high-frequency current steering co-stimulation of subthalamic nucleus and substantia Nigra. Brain Stimul 13, 280–283. 10.1016/j.brs.2019.10.010.

88. Ikeda, H., Kotani, A., Koshikawa, N., and Cools, A.R. (2010). Differential role of GABAA and GABAB receptors in two distinct output stations of the rat striatum: studies on the substantia nigra pars reticulata and the globus pallidus. Neuroscience 167, 31–39. 10.1016/j.neuroscience.2010.01.054.

89. Valldeoriola, F., Munoz, E., Rumia, J., Roldan, P., Camara, A., Compta, Y., Marti, M.J., and Tolosa, E. (2019). Simultaneous low-frequency deep brain stimulation of the substantia nigra pars reticulata and high-frequency stimulation of the subthalamic nucleus to treat levodopa unresponsive freezing of gait in Parkinson’s disease: A pilot study. Parkinsonism Relat Disord 60, 153–157. 10.1016/j.parkreldis.2018.09.008.

90. Weiss, D., Breit, S., Wachter, T., Plewnia, C., Gharabaghi, A., and Kruger, R. (2011). Combined stimulation of the substantia nigra pars reticulata and the subthalamic nucleus is effective in hypokinetic gait disturbance in Parkinson’s disease. J Neurol 258, 1183–1185. 10.1007/s00415-011-5906-3.

91. So, R.Q., McConnell, G.C., August, A.T., and Grill, W.M. (2012). Characterizing effects of subthalamic nucleus deep brain stimulation on methamphetamine-induced circling behavior in hemi-Parkinsonian rats. IEEE Trans Neural Syst Rehabil Eng 20, 626–635. 10.1109/TNSRE.2012.2197761.

92. Chao, T.H., Lee, B., Hsu, L.M., Cerri, D.H., Zhang, W.T., Wang, T.W., Ryali, S., Menon, V., and Shih, Y.I. (2023). Neuronal dynamics of the default mode network and anterior insular cortex: Intrinsic properties and modulation by salient stimuli. Sci Adv 9, eade5732. 10.1126/sciadv.ade5732.

93. Lee, S.H., Broadwater, M.A., Ban, W., Wang, T.W., Kim, H.J., Dumas, J.S., Vetreno, R.P., Herman, M.A., Morrow, A.L., Besheer, J., et al. (2021). An isotropic EPI database and analytical pipelines for rat brain resting-state fMRI. Neuroimage 243, 118541. 10.1016/j.neuroimage.2021.118541.

94. Lee, S.H., Shnitko, T.A., Hsu, L.M., Broadwater, M.A., Sardinas, M., Wang, T.W., Robinson, D.L., Vetreno, R.P., Crews, F.T., and Shih, Y.I. (2023). Acute alcohol induces greater dose-dependent increase in the lateral cortical network functional connectivity in adult than adolescent rats. Addict Neurosci 7. 10.1016/j.addicn.2023.100105.

95. Cox, R.W. (1996). AFNI: software for analysis and visualization of functional magnetic resonance neuroimages. Comput Biomed Res 29, 162–173. 10.1006/cbmr.1996.0014.

96. Yushkevich, P.A., Piven, J., Hazlett, H.C., Smith, R.G., Ho, S., Gee, J.C., and Gerig, G. (2006). User-guided 3D active contour segmentation of anatomical structures: significantly improved efficiency and reliability. Neuroimage 31, 1116–1128. 10.1016/j.neuroimage.2006.01.015.

97. Avants, B.B., Epstein, C.L., Grossman, M., and Gee, J.C. (2008). Symmetric diffeomorphic image registration with cross-correlation: evaluating automated labeling of elderly and neurodegenerative brain. Med Image Anal 12, 26–41. 10.1016/j.media.2007.06.004.

98. Verbeke, G., Molenberghs, G., and Verbeke, G. (1997). Linear mixed models for longitudinal data (Springer).

99. Sobel, M.E. (1986). Some new results on indirect effects and their standard errors in covariance structure models. Sociological methodology 16, 159–186.

100. Paxinos, G., and Watson, C. (2006). The rat brain in stereotaxic coordinates: hard cover edition (Elsevier).

