## Supplemental Figures for "Optogenetic fMRI reveals therapeutic circuits of subthalamic nucleus deep brain stimulation"

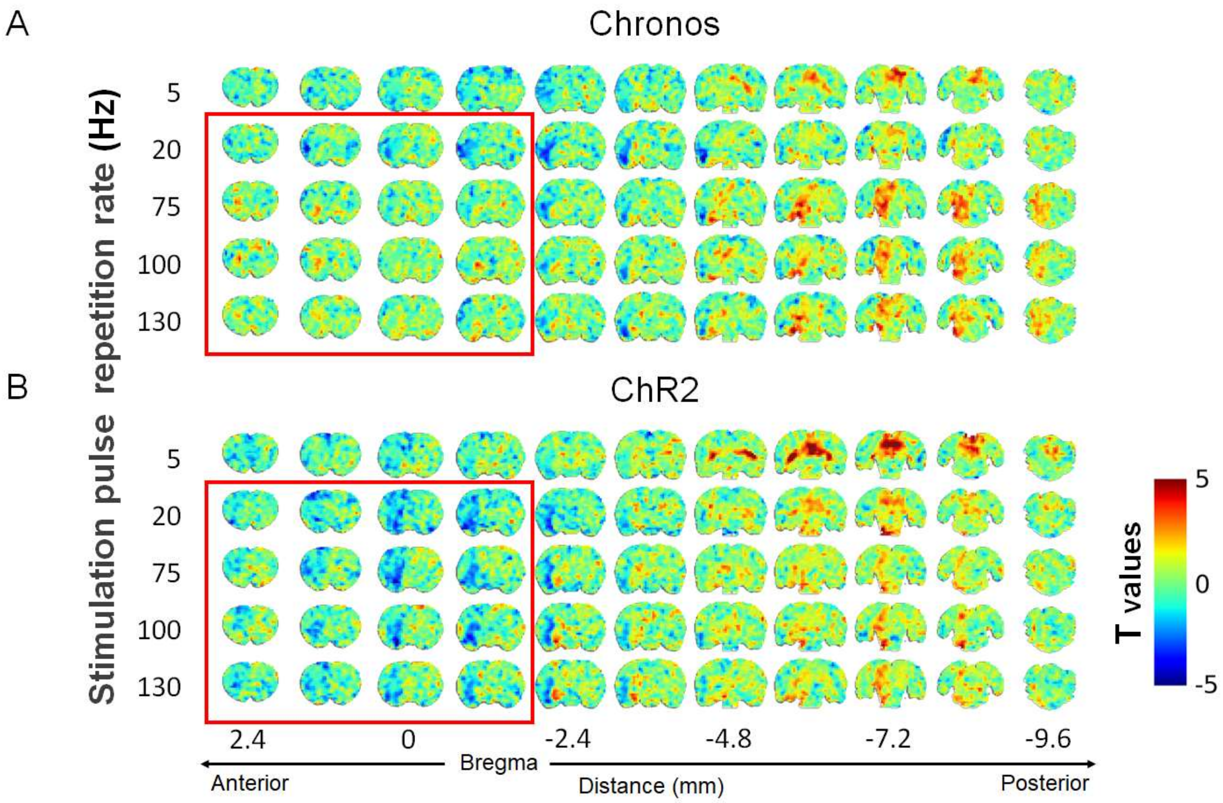


**Fig. S1.**

Group-averaged response maps for each stimulation pulse repetition rate and opsin type. The red box highlights visually identifiable differences between opsin types observed at relatively high pulse repetition rates (above 75 Hz), corresponding to behaviorally effective optogenetic deep brain stimulation (DBS).

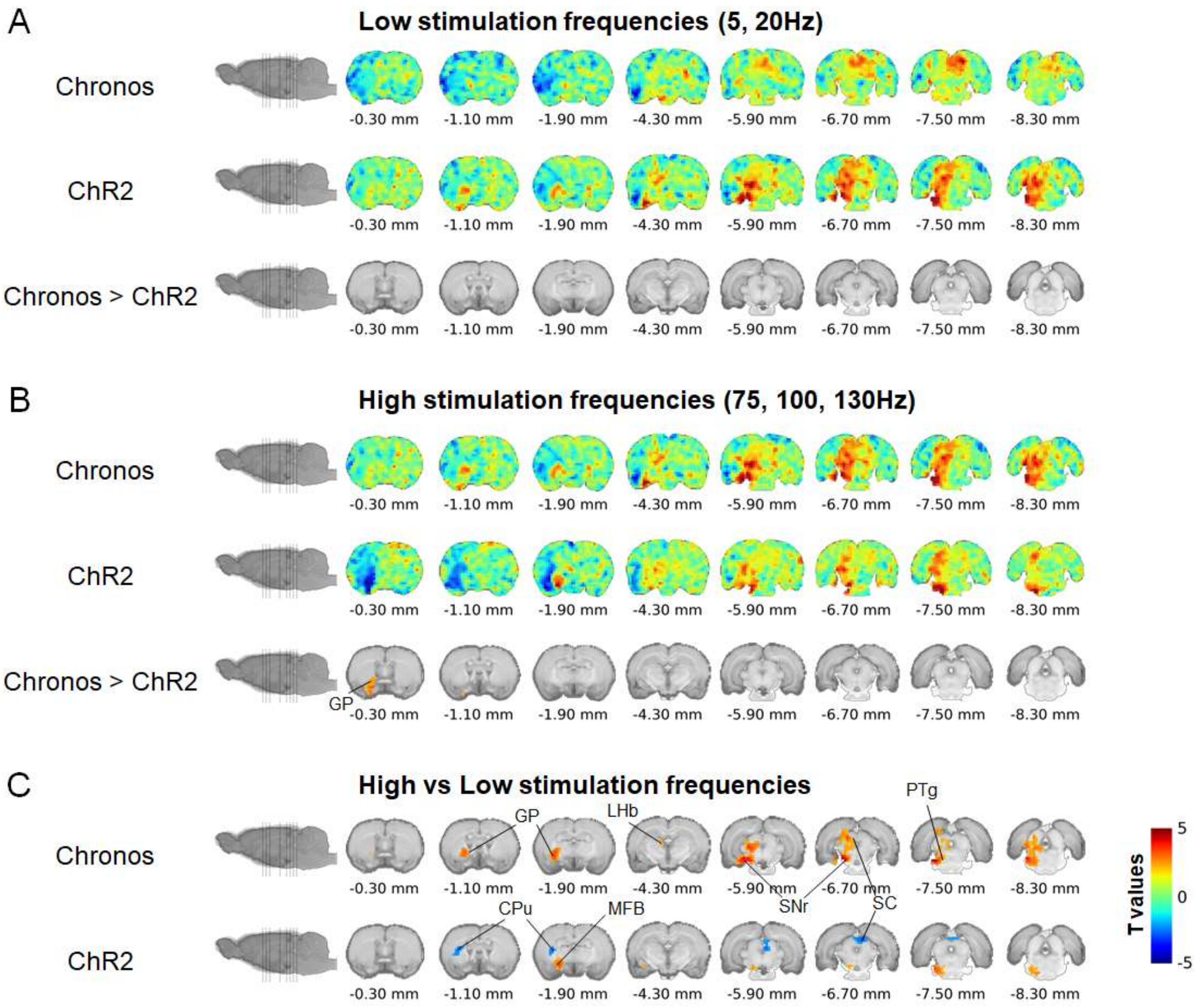


**Fig. S2.**

Averaged contrast maps for low and high stimulation frequencies in each group and their within-group frequency band comparisons. (A, B) Averaged contrast maps representing low and high stimulation frequencies for each group, along with the difference maps between these frequencies. (C) Within-group comparisons highlight the differences between low and high-frequency stimulation. Statistical significance is set at $p < 0.05$, with multiple comparison correction applied.

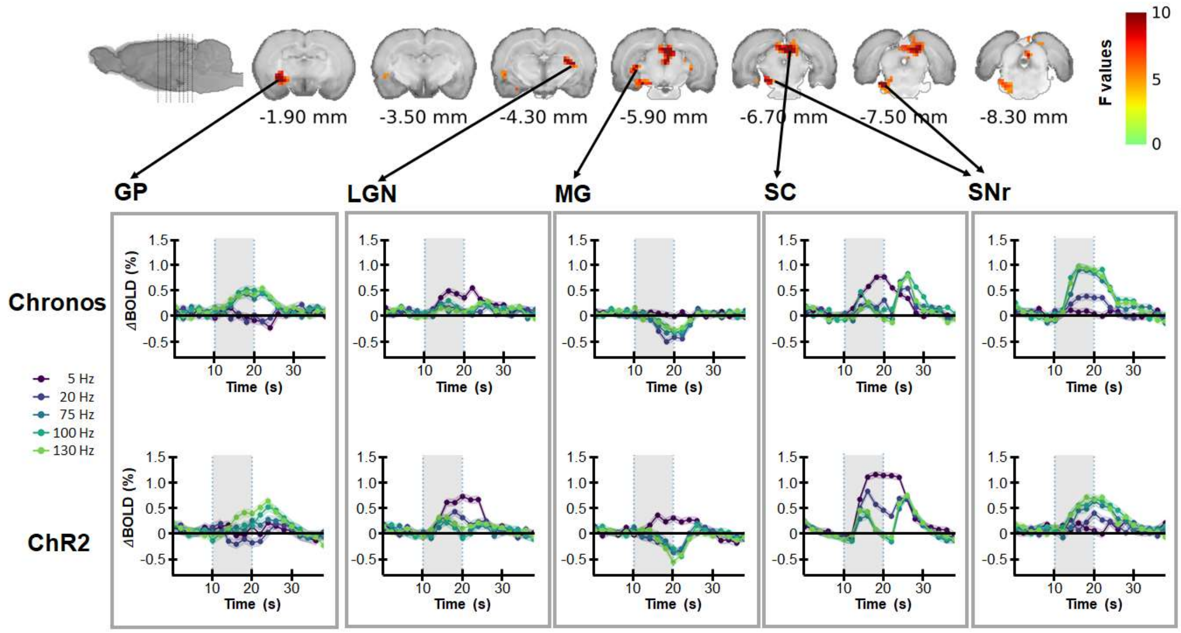


**Fig. S3.**

Statistical parametric maps of significant clusters representing the main effect of stimulation frequency in optogenetic stimulation of the STN for both Chronos and ChR2 groups. Key regions include the globus pallidus (GP), lateral geniculate nucleus (LGN), medial geniculate nucleus (MG), superior colliculus (SC), substantia nigra reticulata (SNr). Statistical significance is set at $p < 0.05$, with multiple comparison correction applied.

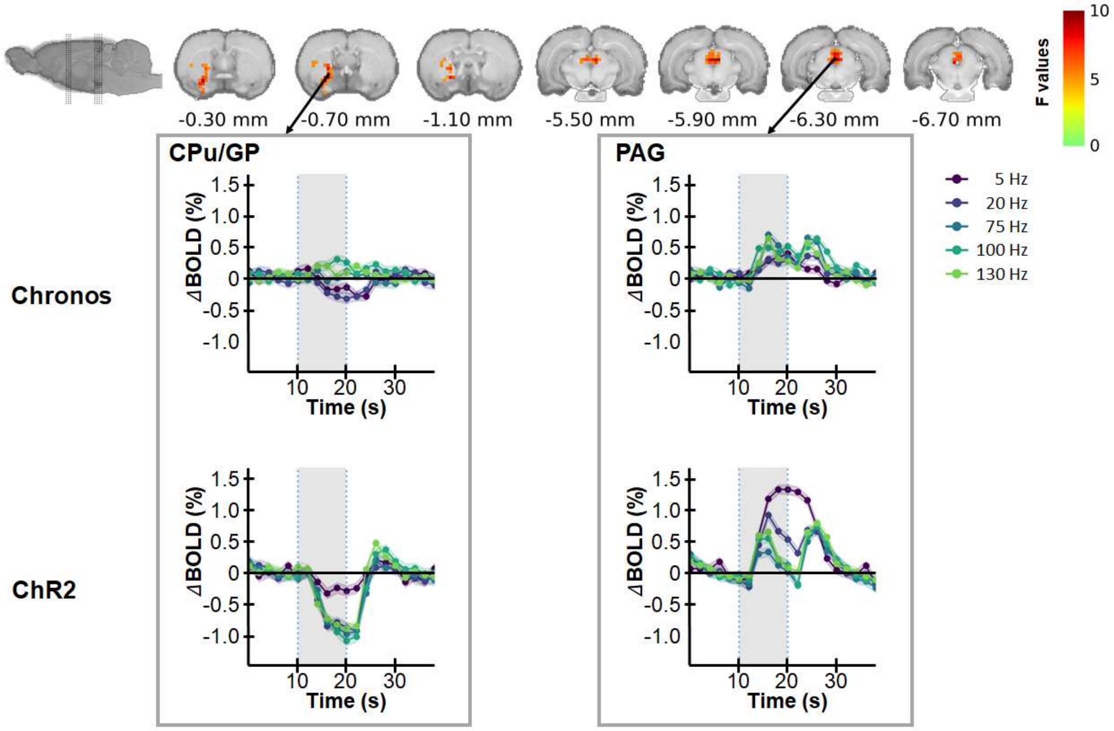


**Fig. S4.**

Statistical parametric maps of significant clusters representing the interaction effect between stimulation frequency and injected virus (Chronos vs $ChR_2$). Key regions include the caudate putamen (Cpu), globus pallidus (GP), and periaqueductal gray (PAG). Statistical significance is set at $p < 0.05$, with multiple comparison correction applied.

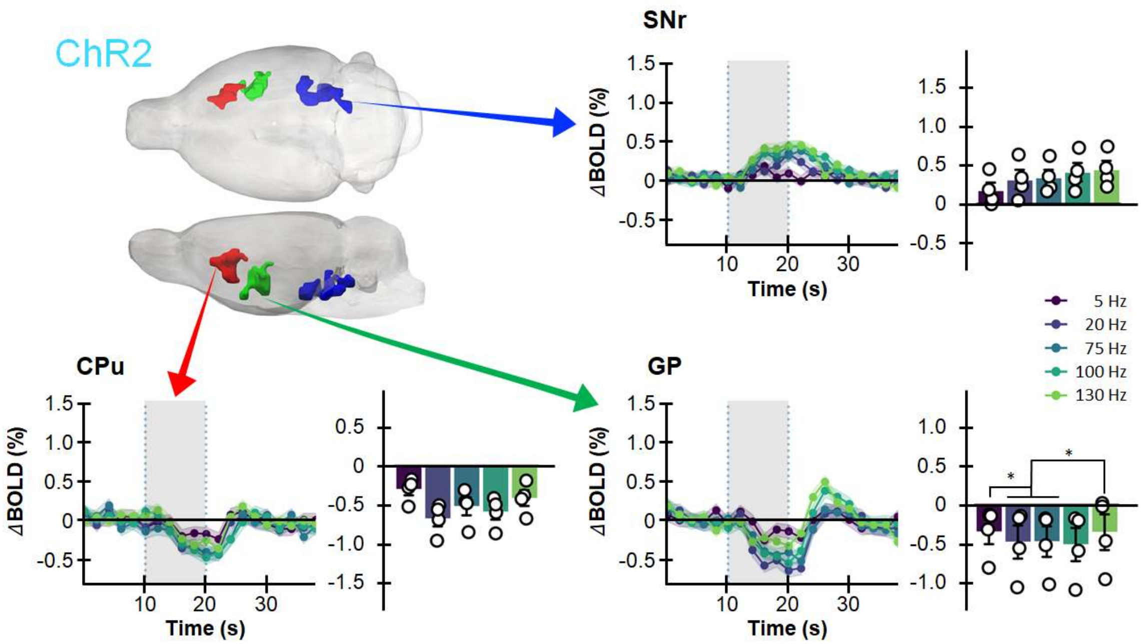


**Fig. S5.**
3D representation of identified ROIs with significant DBS pulse repetition rate effects in the Chronos group and the averaged raw BOLD responses over time for various optogenetic DBS pulse repetition rates at these ROIs in the ChR2 group. Key regions are color-coded as follows: red for the caudate putamen, green for the globus pallidus, and blue for the substantia nigra reticulata. The accompanying bar graphs elucidate the peak of percentage changes of BOLD response, with a T-test used to ascertain significant difference across frequencies (* denotes $p < 0.05$).

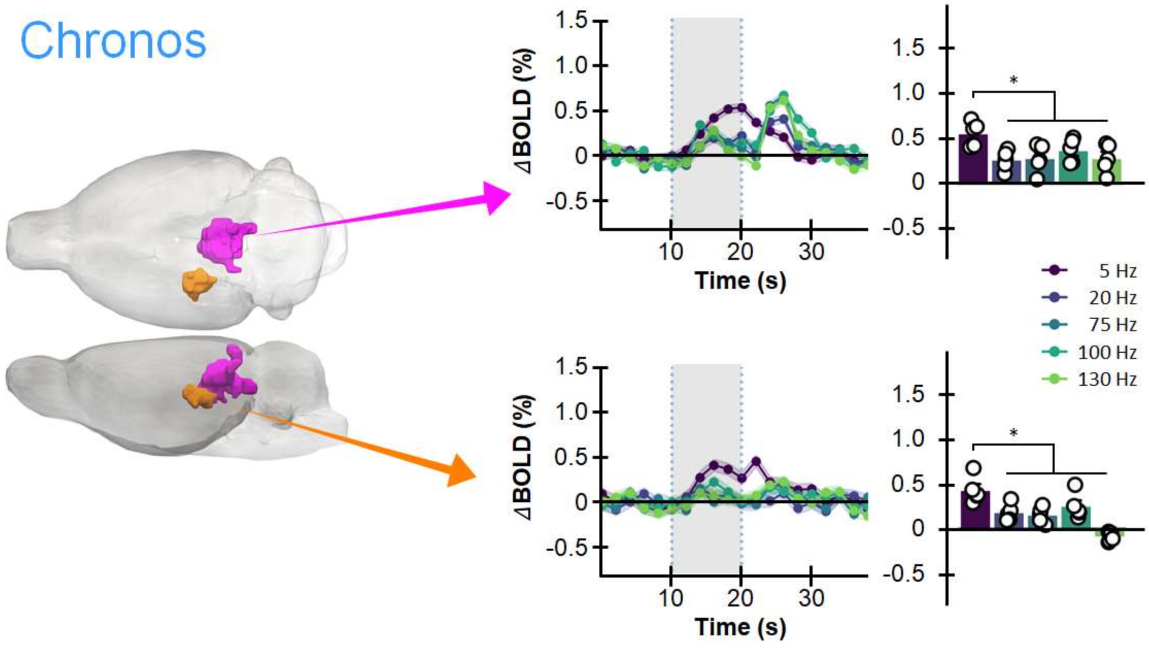


**Fig. S6.**

3D representation of identified ROIs with significant DBS pulse repetition rate effects in the ChR2 group, and the averaged raw BOLD responses over time for various optogenetic DBS pulse repetition rates at these ROIs in the Chronos group. Key regions are color-coded as follows: orange for the dorsal lateral geniculate nucleus, and purple for the superior colliculus. The accompanying bar graphs elucidate the peak of percentage changes of BOLD response, with a T-test used to ascertain significant difference across frequencies (* denotes $p < 0.05$).

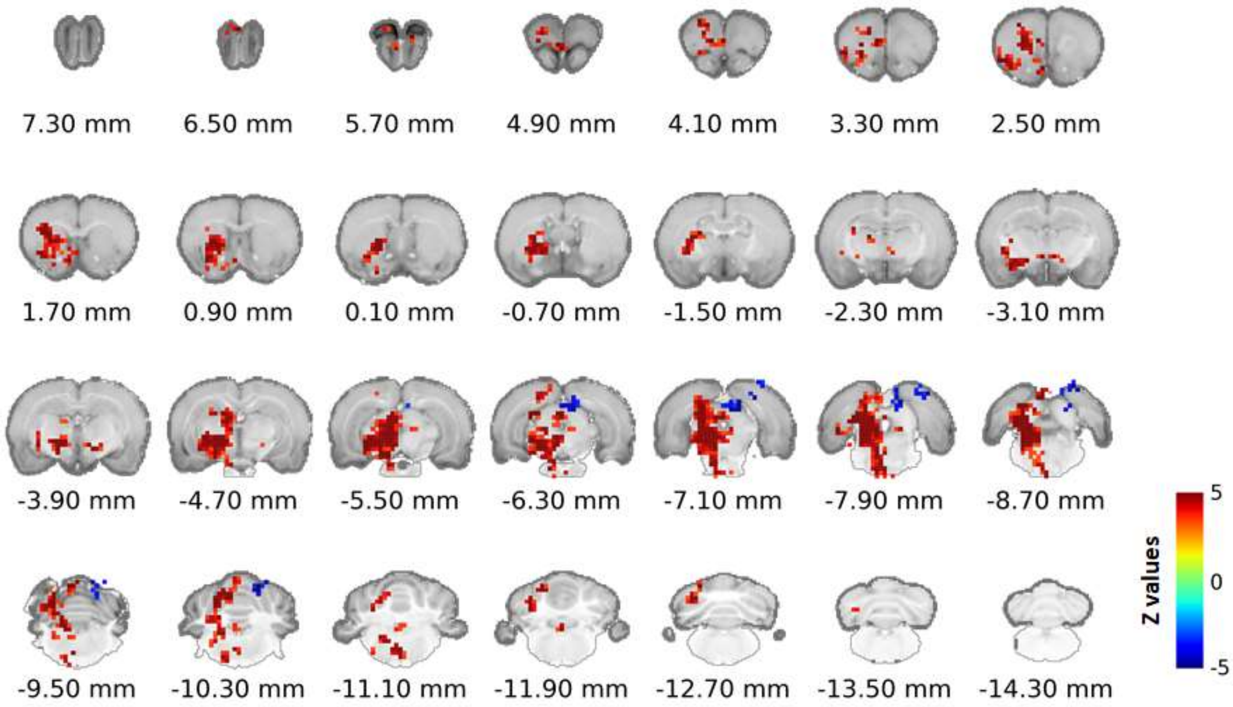


**Fig. S7.**

Color-coded maps exhibit regions where BOLD signal variations, linked to different pulse repetition rates in the Chronos group, correlate significantly with circling behavior (details in Figure 2B). Significance is indicated by a false discovery rate-corrected p-value < 0.001 and a cluster size of more than 40 voxels. The maps are superimposed on coronal brain slices with the left subthalamic nucleus (STN) as the stimulation site. Z scores on the color scale derive from standardized Fisher-transformed Pearson correlations, relating optogenetic STN stimulation response patterns to behavioral data.
